# Population and community responses to the fast, slow, and seasonal components of environmental variation

**DOI:** 10.64898/2026.03.18.712754

**Authors:** Daniel Hernández-Carrasco, Gabrielle Koerich, Anthony J. Gillis, Holly A. L. Harris, Naomi R. Heller, Christina McCabe, Rory S. Lennox, Ilya Shabanov, Li Wang, Hao Ran Lai, Jonathan D. Tonkin

**Affiliations:** School of Biological Sciences, University of Canterbury, Christchurch 8140, New Zealand; Department of Aquatic Ecology, Swiss Federal Institute of Aquatic Science and Technology (Eawag), Dübendorf CH-8600, Switzerland; Oak Ridge Institute for Science and Education, Oak Ridge, Tennessee, USA; School of Biological Sciences, Victoria University of Wellington, Wellington, New Zealand; Bioprotection Aotearoa Centre of Research Excellence, University of Canterbury, Christchurch 8140, New Zealand; South East Asian Rainforest Research Partnership (SEARRP), Kota Kinabalu, Malaysia; Te Pūnaha Matatini Centre of Research Excellence, University of Canterbury, Christchurch 8140, New Zealand

**Author notes:** **Corresponding authors:** D.H.-C, G.K. These authors contributed equally as joint first authors (Daniel Hernández-Carrasco, Gabrielle Koerich).

## Abstract

Theory suggests that different components of environmental fluctuations, from daily and seasonal cycles to multidecadal trends, can have distinct and even opposing effects on species’ abundances and community dynamics, depending on their specific adaptations. But empirical research that deconstructs the influence of these different cycles on communities is lacking. Here, we used long-term biological monitoring data together with flow records of rivers across New Zealand to (i) investigate the role of fast, slow, and seasonal river-flow fluctuations in structuring macroinvertebrate communities; and (ii) to assess whether life-history and mobility traits mediate the response. Using joint species distribution models, we found striking differences in taxon and community responses to the different components of river flow variation. Responses to slow fluctuations were generally stronger and better predicted by traits, while responses to seasonal fluctuations were highly heterogeneous. Fast increases in flow, typical of flooding events, had pervasive negative effects on species abundances, but the severity of impact partly depended on mobility traits. Our results suggest that different ecological mechanisms underpin the response to distinct environmental fluctuations, highlighting the value of jointly considering multiple temporal scales of variation and species’ functional traits to understand and predict how communities reorganise under fluctuating environmental regimes.

## 1 Introduction

Environmental fluctuations are a fundamental driver of temporal changes in biodiversity. These fluctuations can exhibit complex temporal patterns, arising from multiple sources of variation operating at different characteristic frequencies (Wolkovich et al., 2014). For instance, most ecosystems are subject to daily and seasonal thermal cycles superimposed on longer-term nonstationary trends such as multidecadal warming or cooling. Theory suggests that these different components of environmental variation can strongly shape ecological and evolutionary responses (Hernández-Carrasco et al., 2025; Pande et al., 2020; Picoche & Barraquand, 2019; Ruokolainen et al., 2009; Sakavara et al., 2018; Yang et al., 2019) but attempts to incorporate them in empirical analyses remain relatively scarce (but see Brodie et al., 2021). This gap is particularly critical given that ongoing climatic changes, including directional long-term trends and increasing frequency and intensity of extreme events, are intrinsically tied to distinct components of temporal environmental variation (Chen et al., 2025; Harris et al., 2018; Tonkin et al., 2026; Walker & Van Loon, 2023; Wang et al., 2024).

Population- and community-level responses to environmental fluctuations likely depend on their frequency relative to organisms’ life cycles, as it determines which individual- and population-level mechanisms are at play (Bieg et al., 2023). When fluctuations occur within an organism’s lifespan, populations can present dampened demographic responses due to physiological and population-level processes that integrate environmental conditions, including energy storage, lagged vital rates, dispersal, and diverse bet-hedging life-history strategies (Heino et al., 2000; Mustin et al., 2013; Ripa & Lundberg, 1996; Varpe, 2017). Such mechanisms have evolved because high temporal variability in population growth rates (i.e., responding strongly to temporal environmental fluctuations) typically reduces long-term growth and increases extinction risk (Hernández-Carrasco et al., 2025; Hilde et al., 2020; Starrfelt & Kokko, 2012). Conversely, theoretical models suggest that populations with specific life-history traits such as high and density-dependent fecundity and short generation times can present strong responses to faster fluctuations (Grosholz et al., 2021; Koons et al., 2009; Ruokolainen et al., 2009).

Prolonged environmental changes, particularly those with a frequency lower than organisms’ lifespans, could have elevated impacts on populations and communities compared to faster fluctuations for a variety of reasons. Potential mechanisms involved in these responses include the physiological deterioration of organisms under extended suboptimal conditions (Wingfield, 2013), higher risk of local extirpation during periods of low abundance (Mustin et al., 2013), and a greater influence of antagonistic species interactions (Gudmundson et al., 2015; Ruokolainen & Fowler, 2008). However, slow environmental fluctuations may also provide enough time for behavioural and plastic responses to stabilise populations that possess such adaptations (Bernhardt et al., 2020; Postuma et al., 2020). For species whose optimum conditions differ from the average environment, slow environmental transitions towards relatively more favourable states imply extended opportunities for local colonisation and greater increases in abundance during those periods (Gonzalez & Holt, 2002; Picoche & Barraquand, 2019). Thus, slow fluctuations could promote community change through a combination of positive and negative species responses to the environment (balanced abundance changes), resulting in the replacement of species by those better adapted to the transient environment (Stemkovski et al., 2025).

In addition to frequency, periodicity can be important in determining ecological responses to environmental variation (Hernández-Carrasco et al., 2025). Periodic fluctuations, such as daily or annual rhythms, allow organisms to synchronise their life cycles and behaviour with predictable environmental conditions within the day or year (Varpe, 2017). For instance, many insects, birds, and mammals synchronise long-distance migrations with predictable seasonal patterns (Bauer & Hoye, 2014; Satterfield et al., 2020). These adaptations can lead to strong changes in population abundances and seasonal turnover of species composition, as organisms may be active during favourable periods and inactive during unfavourable ones (e.g., through movement patterns, dormant forms, and life cycles with distinct realm-specific stages) (Humphries et al., 2017; McMeans et al., 2015). However, when seasonal fluctuations are not the predominant signal, strategies to cope with unpredictable sources of environmental variation can reduce population- and community-level responses to seasonal fluctuations (Tonkin et al., 2017). A challenge, therefore, lies in understanding the complex ecological responses to multi-scale temporal environmental fluctuations inherent to dynamic systems, such as rivers.

Rivers exhibit a wide range of co-occurring fluctuations in discharge (hereafter ‘flow’), which is a key driver of population and community dynamics (Palmer & Ruhi, 2019; Poff et al., 1997). River flow can vary seasonally due to predictable changes in snow melt, evapotranspiration, and rainfall; over weeks to months due to sudden rainfall events or short periods of extreme evapotranspiration; and over years to decades due to climatic drivers such as El Niño–Southern Oscillation or anthropogenic climate change (Wang et al., 2024; Ward et al., 2010; Yuan et al., 2023). Given natural flow regimes encompass all of these different frequencies (Tonkin et al., 2021), rivers are ideal systems to assess ecological responses to the various components of environmental variation. Among riverine communities, benthic macroinvertebrates comprise species with diverse life histories and mobility traits that could determine their responses to different scales of temporal environmental fluctuations (Lytle & Poff, 2004). For instance, macroinvertebrate species can range from those with multiple generations per year through to multi-year life cycles, and those with fully aquatic life cycles through to species with a terrestrial adult phase (Verberk et al., 2008). This trait variability could provide different degrees of sensitivity to environmental fluctuations.

Here, we aimed to disentangle species and community-level responses to fast (weeks to months), slow (months to years), and seasonal fluctuations in river flow using a long-term monitoring dataset of macroinvertebrate communities across New Zealand. We jointly modelled macroinvertebrate species responses to decomposed flow variation and assessed the relationship between environmental responses and life-history and mobility traits. We considered four competing scenarios for how macroinvertebrate communities might respond to different flow components:

i. Homogeneous response: Environmental fluctuations exert uniform effects across taxa and temporal scales. In this null scenario, neither species identity nor the frequency of fluctuations would moderate changes in abundances, implying a lack of functional differentiation in flow responses.
ii. Frequency-dependent, taxon-invariant response: Different flow components have distinct community-level effects on taxa abundances, but taxa respond uniformly within each component. This would suggest disparate ecological mechanisms are engaged at different frequencies of flow variation, but those mechanisms would be independent of taxa identity and their trait variability.
iii. Frequency-independent, taxon-specific response: Flow has distinct effects on taxa abundances that depend on their identity but not on the frequency of fluctuations. Taxa responses are therefore positively correlated across flow components. This scenario would imply that the mechanisms involved in the response to flow vary across species, but are homogeneous across flow frequencies.
iv. Frequency-dependent, taxon-specific response: Different flow components exert distinct effects on abundance that also vary among taxa. The correlation between taxon-specific responses to different flow components is therefore low. This scenario would suggest different mechanisms involved in the response to flow that depend on both frequency and species characteristics.

We expected macroinvertebrate responses to be most consistent with scenario (iv) as different flow components likely engage distinct ecological mechanisms that also depend on species functional characteristics. We expected taxa responses to be stronger for slow fluctuations (due to prolonged stress, growth, and colonisation opportunities) and seasonal fluctuations (due to phenological synchronisation), and weaker for fast fluctuations (due to dampening mechanisms and adaptations to naturally dynamic environments; Winterbourn et al., 1981). At the community level, we expected slow and seasonal components to drive balanced abundance changes—turnover in the abundance of different taxa—reflecting the ability of some populations to track these environmental fluctuations with increases in abundance. Our study constitutes one of the first attempts at empirically testing community responses to multiple components of temporal environmental variation.

## 2 Methods

### 2.1 Study location

Aotearoa New Zealand is a narrow oceanic island nation that supports a diverse range of landscapes over a relatively large latitudinal gradient (35*^◦^*S to 46*^◦^*S). Consistent with its oceanic climate, rainfall events and subsequent floods tend to be frequent and relatively unpredictable, resulting in fast changes in river flow throughout typically short river continua (Winterbourn et al., 1981). Despite this, New Zealand rivers also undergo slower, interannual fluctuations associated with atmospheric circulation patterns, such as the El Niño Southern Oscillation (ENSO) and the Southern Annual Mode (SAM) (Chiew & McMahon, 2002; Li & McGregor, 2017), and periodic annual variability due to snowmelt and rainfall seasonality (Queen et al., 2023)—albeit much less predictable than continental systems where snowmelt dominates the discharge signal. Thus, river flow regimes across New Zealand comprise slow, fast, and seasonal components of variation.

### 2.2 Macroinvertebrate data

We retrieved macroinvertebrate abundance data from the New Zealand National River Water Quality Network (NRWQN), maintained by the Earth Sciences New Zealand (formerly, NIWA). The monitoring scheme covers 60 sites on 42 rivers sampled on an annual basis between 1990 and 2020 (Fig. S1), mostly during the late austral summer (84.8% of samples), and selected for their accessibility, proximity to hydrological recording stations, and national or scientific importance (Smith & McBride, 1990). In addition, sites were selected to represent different degrees of human impact within each region. Sites are located between latitudes 46.38*^◦^*S and 35.27*^◦^*S, covering most of New Zealand’s North and South Islands, and at altitudes ranging between 2 and 655 metres above sea level (median: 111 m asl). Stream orders vary between 3 and 8 (median: 6).

Seven 0.1 *m*^2^ Surber samples were collected over as many substrate types as possible, and individuals were counted and identified to the lowest practical taxonomic unit (Smith & McBride, 1990). We kept taxa that occurred in at least 50 samples (3.14% of samples), and at least 5 unique sites (8.33% of sites). Because the taxonomic resolution has improved during the course of the monitoring program, we manually matched taxon names to ensure agreement with the highest taxonomic resolution that was consistent throughout the dataset. We were left with 67 individual taxa corresponding to either Genus (50), Tribe (2), Subfamily (3), Family (9), Order (1; Amphipoda), Subclass (2; Acarina and Hirudinea), Class (2; Oligochaeta and Ostracoda) or Phylum (2; Nematoda and Platyhelminthes).

We retrieved life-history and dispersal trait data from the New Zealand Freshwater Macroinvertebrate Trait Database (Phillips & Smith, 2018). Focusing on traits related to frequencies of temporal environmental variation, we selected five life-history and dispersal traits: (1) dissemination potential, (2) aquatic stages (i.e., which stages within the life cycle are aquatic), (3) aquatic mobility, (4) number of reproductive cycles per year, and (5) adult body size. We transformed ‘fuzzy-coded’ trait scores to either weighted means for continuous traits (e.g., adult body size) or normalised categories for discrete traits (e.g., aquatic stages). In cases where trait data had a higher taxonomic resolution than the abundance data after processing (16% of taxon groups), we used the average trait values. In addition to traits, we retrieved taxon sensitivity scores used for the computation of macroinvertebrate-based water quality indices (Stark & Maxted, 2017) to assess whether our models had successfully accounted for known patterns in community composition along gradients of human impact and river size.

### 2.3 Environmental data

#### 2.3.1 Flow decomposition

We sourced daily means of river discharge for sites associated with abundance data from 1982 to 2022 from Earth Sciences New Zealand Hydro Web Portal (hydrowebportal.niwa.co.nz). To capture flow variability at multiple temporal scales, we decomposed log-river discharge into mean, seasonal, slow, and fast components of temporal variation. First, we extracted the seasonal component by fitting a linear combination of Fourier basis (periodic at the annual frequency) to each flow time series. We used 10 basis functions, which was sufficient to capture relatively complex annual patterns. We then applied two consecutive convolutions to the seasonally de-trended flow data using half-Gaussian kernels to obtain the slow and fast components. This procedure effectively computes weighted moving averages with weights determined by the shape of the kernel. Thus, fast and slow flow components depend only on previous time points, with a greater influence of recent flow values. We used kernels with a standard deviation of 365 days (∼95% of weight within the previous two years) and 45 days (∼95% of weight within the previous three months) to obtain the slow and fast components of flow, respectively. Gaps in flow data of up to 14 days were imputed using the na_seadec function in the *imputeTS* R package (Moritz & Bartz-Beielstein, 2017). The seasonal component was computed using the seasonal.smooth function in the *mar1s* R package (Paramonov, 2018).

To compare the magnitude of the *k* = 3 flow components (*c*) in each river’s flow regime (*i*), we calculated their contribution to the observed variance,

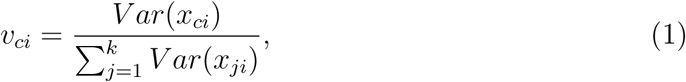

where *v_ci_* reflects the proportion of the variability in log-flow of stream *i* attributed to the flow component *c*.

#### 2.3.2 Human Impact Index

To account for variation in the degree of anthropogenic impact within our dataset, we used the average Stock Unit Density (SUD) of cattle, sheep, and deer in the catchment area above each sampling point. Land use, and particularly the intensity of farming, is strongly related to water quality indicators such as nitrogen and phosphorus concentration in the sampled streams (Julian et al., 2017), making it a good proxy of anthropogenic impact. To make the SUD of different animals comparable, we computed Total Nitrogen Excretion Rates (TNER) based on animal-specific annual excretion rates (Luo & Kelliher, 2010) and aggregated them for each catchment. SUD values were interpolated between two available years (1990 and 2011) or assigned the closest value. We included TNER in subsequent analyses. Stock Unit Density data were obtained from Julian et al. (2017).

### 2.4 Community modelling framework

We quantified taxon-specific abundance responses to each component of flow variation using Generalised Linear Latent Variable Models (GLLVM; Niku et al., 2019, 2025). GLLVMs expand generalised linear mixed-effects models by including a small number of latent variables with species-specific loadings to reduce the dimensionality of random effects in the model, and account for potential residual correlations between species abundances. We included site as a random effect determining the position of samples within two latent variables, and a random effect of time with an exponentially decaying covariance structure to account for temporal autocorrelation. We specified a negative binomial distribution and the canonical log-link function to account for the discrete nature of count data and potential overdispersion.

To quantify the role of life-history and dispersal traits in modulating responses to different components of flow fluctuations, we also fitted a fourth-corner model (Brown et al., 2014). For ‘fuzzy-coded’ traits (i.e., those coded as a degree of association to each trait modality), we used the most common modality as a baseline and removed it from the trait matrix **T** (taxa × traits) to avoid perfect collinearity issues. Therefore, the slopes of the baseline trait modalities were captured by the main environmental effects (***β****_e_*). In addition to mediation by traits, we allowed for taxon-specific responses not strictly determined by traits (***β****_j_*) *via* random taxon slopes for each flow component (**z***_j_*):

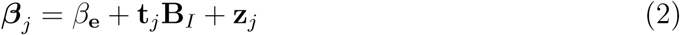

where **t***_j_* ∈ **T** is a vector of traits of taxon *j* and **B***_I_* is a matrix of associated fourth-corner coefficients (traits × environment). See Niku et al. (2019) for further details on the full GLLVM fourth-corner model parametrisation.

We quantified the overall association between traits and taxon-specific responses to each flow component (*c*) by computing a Pseudo-*R*^2^ akin to Nakagawa and Schielzeth (2013):

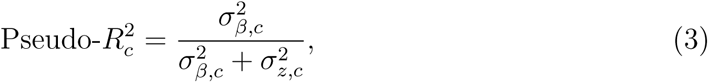

where 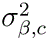 is the variance of marginal slopes for flow component *c*, and 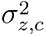 is the variance associated with taxon-specific random slopes. Values of 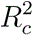 close to one indicate that variation in taxon responses to the flow component *c* (not taxon abundances) can be perfectly explained by the traits included in the model, whereas values closer to zero indicate that most of the variability is not related to traits.

We further analysed trait-mediated environmental effects by computing marginal slopes for counterfactual trait combinations. Given the complexity of the fourth-corner model, robust confidence intervals for trait-mediated responses were computed using single-fit parametric bootstrapping instead of the delta method (Fletcher & Jowett, 2022). In brief, parametric bootstrapping consists in drawing samples from a multivariate normal distribution MVN(*θ,* **Σ**) defined by the vector of estimated model parameters (*θ*) and their variance-covariance matrix (**Σ**). Because GLLVM can be sensitive to starting parameter values, we fitted ten versions of each model with small, random starting values and selected the model with the highest log-likelihood (Niku et al., 2019).

### 2.5 Analysing community-level dynamics

To assess community-level responses to different components of flow fluctuation, we (1) used fitted GLLVMs to obtain expected species abundances at different levels of each flow component and (2) computed pairwise dissimilarities between expected communities. We used the Bray-Curtis dissimilarity index and its two partitions: balanced abundance changes and abundance gradients, which are analogous to the turnover and richness difference components of presence-absence-based dissimilarity (Baselga, 2017). Dissimilarity indices were computed with the R package *betapart* (Baselga et al., 2022). Expected abundances were log-transformed before computing dissimilarities to reduce the disproportionate influence of a few dominant taxa.

Because spatial variability in community composition can lead to different community-level responses across sites, we repeated the calculations for each river using the loadings in the two latent variables, the river’s mean log-flow, and the river’s mean TNER. In other words, we repeated the calculations for each river accounting for its expected baseline community composition.

## 3 Results

### 3.1 Patterns in flow components across New Zealand

The seasonal pattern of flow variation was relatively homogeneous across our study sites, with most streams presenting higher flow in winter and lower flow at the end of the summer (Fig. 1). Seasonal and fast fluctuations were the main source of log-flow variation across the 60 sites and accounted for an average of 47% and 43% of the observed variability, respectively (Fig. 1C). By contrast, slow fluctuations only accounted for an average of 10% of the total variation in log-flow. Mean stream flow was strongly correlated with river order (Pearson’s correlation of 0.828, Fig. S2), indicating that the average flow captured variability in river size associated with its position in the river network.

**Fig. 1:**
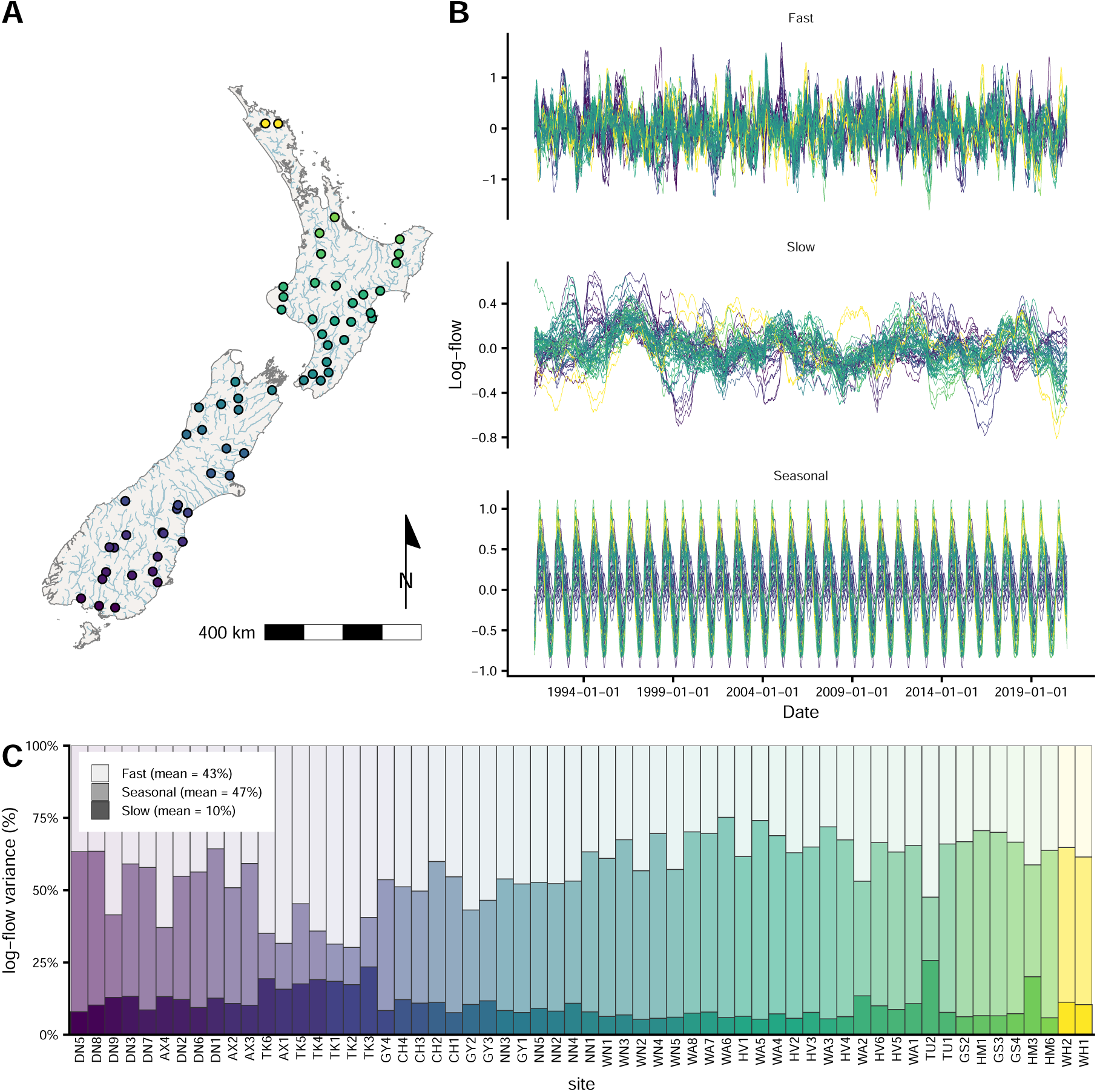
River-flow components across Aotearoa New Zealand. (A) Study sites were located across the North (Te Ika-a-Māui) and South islands (Te Waipounamu). Colour indicates site’s latitude. (B) Values of different river flow components during the study period. (C) Proportion of variance in log-flow explained by each flow component. Each column represents a unique location.

### 3.2 Taxon-specific responses to the components of flow variation

Taxon-specific responses were strongest to slow fluctuations in log-flow compared to other flow components (Fig. 2). Crustacea, Gastropoda, worm-like taxa, and many Trichoptera were associated with prolonged periods of low flow, whereas the responses of Ephemeroptera, Plecoptera, Diptera, and other groups were more diverse. Responses to the fast flow component were mostly negative, with 30 taxon groups showing significantly negative coefficients and only one relatively rare taxon (*Megaleptoperla spp.*) showing a significantly positive response (Fig. 2). *Sigara* spp., Chironominae, and Tanypodidae showed the strongest negative responses to fast increases in log-flow. Taxon-specific responses to seasonal fluctuations were highly diverse across taxonomic groups. Moreover, responses to different flow components were only weakly correlated, with a maximum Pearson’s correlation of 0.233 between responses to seasonal and fast flow fluctuations (Fig. S3).

**Fig. 2:**
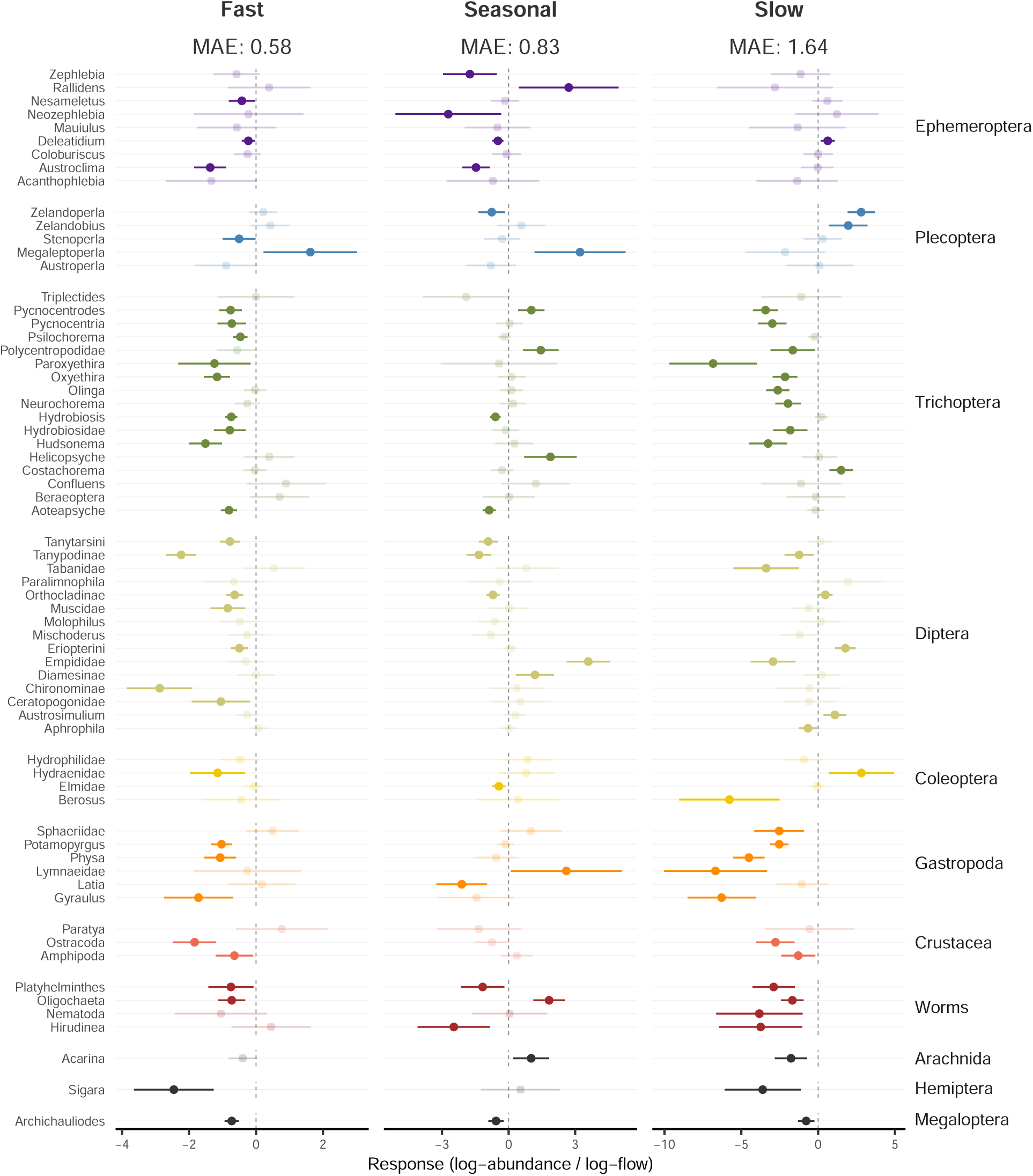
Taxon responses to fast, seasonal, and slow flow fluctuations. Points show estimated taxon-specific effects, while horizontal lines indicate 95% confidence intervals. Colours correspond to different taxon groups as indicated on the right. Light colours indicate cases where 95% CI crosses 0. The mean absolute effect (mean abs) across taxa is indicated for each flow component.

In addition to the different components of flow variation, responses to mean flow and TNER, and sites loadings into the two latent variables suggest that our models successfully accounted for known gradients in community composition driven by land use and water quality (see Supplementary Information S4).

### 3.3 Trait mediation

The fourth-corner model detected significant associations between taxon responses to flow components and their life-history and mobility traits (Fig. 3). Aquatic stages within the life cycle were the most relevant trait determining the response to slow fluctuations. Organisms with a terrestrial phase showed a more positive response to slow increases in flow, and the opposite was true for strictly aquatic organisms (i.e., organisms with aquatic adult and larvae). Although swimming taxa showed only marginally significant effects for the response to fast fluctuations (Fig. 3A), they were the trait modality with the strongest negative response to fast increases in flow (Fig. 3B). Neither the number of reproductive cycles per year nor adult size showed significant effects on the response to fast, seasonal, or slow flow components. Pseudo-*R*^2^ for taxa responses revealed that traits were most relevant for determining the response to the slow flow component 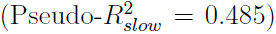 compared to the fast 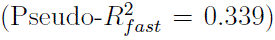 and seasonal components 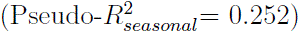 (Fig. 3C).

**Fig. 3:**
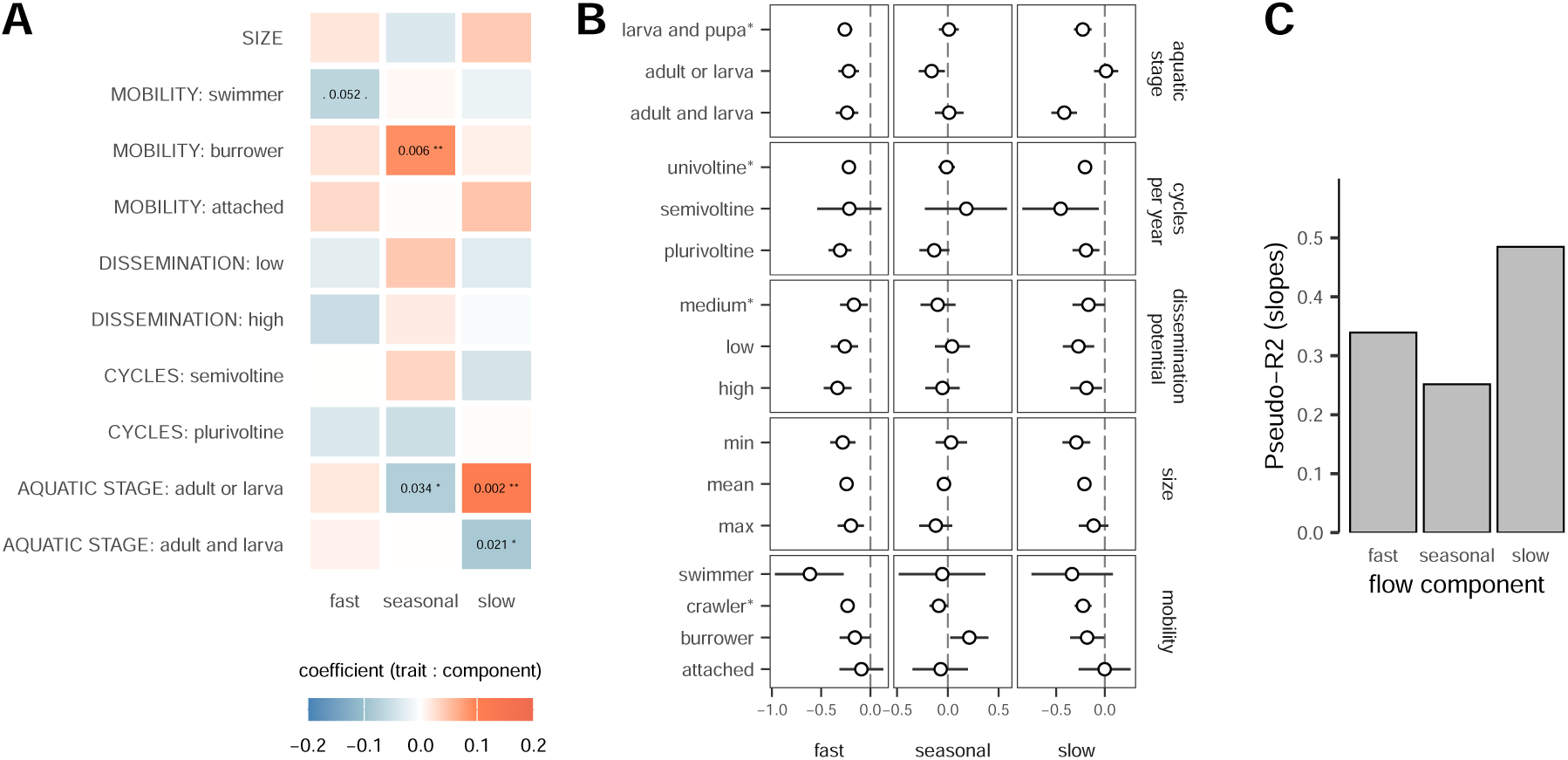
Trait-mediated responses to fast, seasonal, and slow flow fluctuations. (A) Fourth-corner coefficients. Associated *p*-values lower than 0.1 are shown together with standard significance codes (’***’: *<*0.001, ‘**’: *<*0.01, ‘*’: *<*0.05, ‘.’: *<*0.1). (B) Expected slopes for different trait modalities. Horizontal lines indicate 95% credible intervals obtained with single-fit parametric bootstrapping (details in main text). For categorical variables, the modality used as a baseline in the model is indicated (*). (C) Pseudo-*R*^2^ indicating the proportion of variance in taxon-specific responses to flow components (slopes) determined by traits.

### 3.4 Community-level responses

Expected community change based on model predictions showed distinct community-level responses to the different components of flow variation (Fig. 4). Rates of community change (Bray-Curtis per unit of log-flow) across sites were highest for slow fluctuations. Although expected rates of overall community change due to fast and seasonal fluctuations were similar in magnitude, community change in response to fast fluctuations was mostly driven by gradients in taxa abundances, while for seasonal fluctuations community change was almost exclusively driven by balanced abundance changes (Fig. 4).

**Fig. 4:**
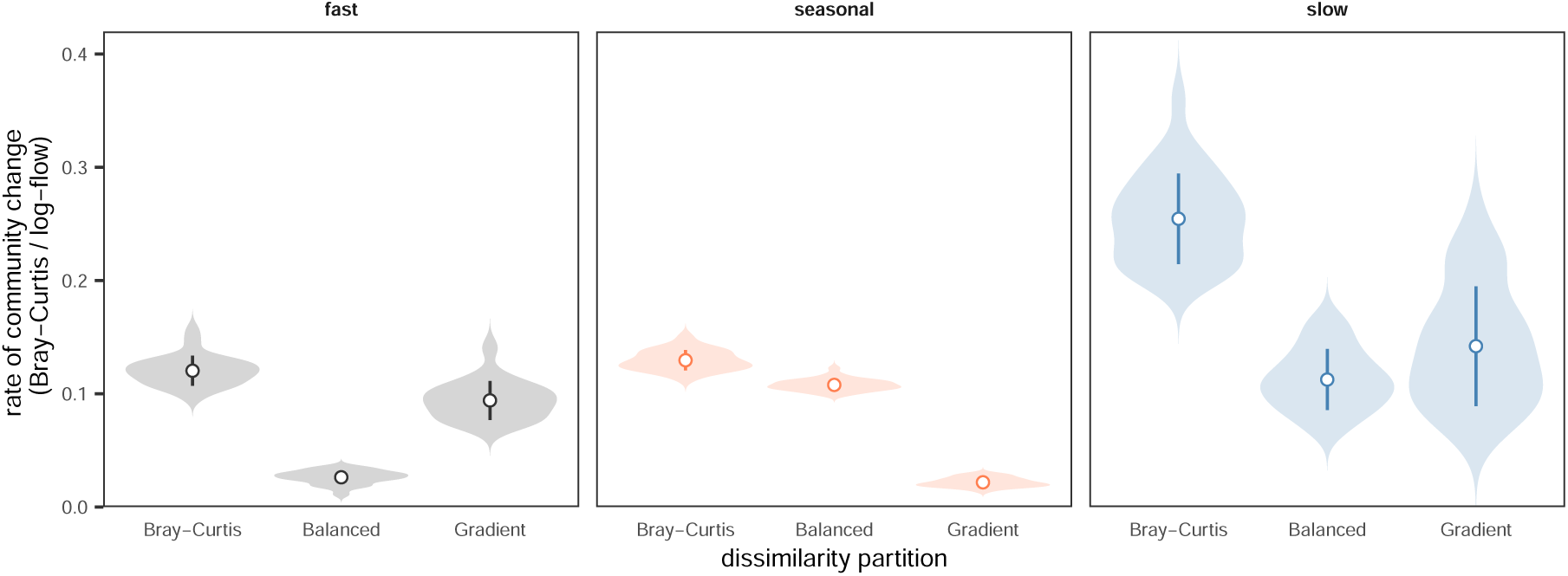
Expected rate of community change along components of flow variation. Circles and line ranges indicate the mean and standard errors across sites. Shaded areas represent the distribution of expected values. Community change was calculated as the Bray-Curtis dissimilarity, and its partitions: Balanced variation in abundance (Balanced) and abundance gradients (Gradient).

## 4 Discussion

Environmental fluctuations occur across a wide gradient of frequencies, from daily to decadal (Wolkovich et al., 2014), shaping the structure of populations and communities across temporal scales (Bernhardt et al., 2018; Duffy et al., 2022). By partitioning river flow regimes into distinct temporal components (slow, fast, and seasonal fluctuations), our study revealed that macroinvertebrate taxon responses were largely uncorrelated across flow components and showed distinct trait associations—consistent with our scenario (iv). This pattern suggests that the mechanisms driving community composition under environmental (flow) fluctuations are specific to the frequencies of such fluctuations, and depend on species functional characteristics. Overall, our findings demonstrate that treating the different components of environmental variation as unique ecological drivers is essential for capturing the complexity of community assembly in dynamic environments.

Despite explaining an average of approximately 10% of the variance in river flow, slow fluctuations generated strong ecological responses. Persistent flow fluctuations, captured by the slow flow component in our analyses, were comparable in their influence on species abundance to larger but faster fluctuations. This pattern aligns with population theory, which suggests that positively autocorrelated environmental fluctuations (i.e., slower fluctuations) have disproportionately large effects on population dynamics and extinction risk (Bernhardt et al., 2020). Empirical studies have similarly shown community restructuring in response to long-term environmental change (Haase et al., 2023; Pinsky et al., 2025). Persistent suboptimal conditions, such as prolonged drought or low-flow conditions, can accumulate into negative carry-over effects that reduce long-term fitness and hinder short-term recovery, potentially leading to pronounced population declines and local extirpations (Giménez et al., 2022). Conversely, extended periods of relatively favourable conditions can provide sufficient time for population growth, allowing populations to track environmental changes through abundance fluctuations. Our results agree with this expectation, showing a higher degree of balanced abundance changes at the community level in response to slow versus fast fluctuations—indicative of deterministic turnover among species with differential adaptation to transient environmental conditions. Yet, community-level changes in response to slow fluctuations were also driven by abundance gradients, with many taxa declining at higher flow levels. Such predominance of negative responses to the slow-flow component implies that mean flow conditions are already near or exceed optimal ranges for many taxa, such that further increases in flow represent a transition towards harsher conditions and therefore abundance declines. Similar patterns of decreasing species abundances and richness have been reported for communities under harsh environmental gradients such as increasing latitudes in communities near the poles (Soininen et al., 2018), and increasing water temperatures in tropical regions (Meyer et al., 2024).

Strong responses to slow fluctuations reflected species-specific sensitivity to prolonged flow conditions. In fact, life-history traits were best at predicting responses to slow flow components compared to fast and seasonal. Taxa with strictly aquatic life cycles tended to be more negatively associated with increasing slow fluctuations, whereas taxa with only one aquatic stage responded more positively. These relationships likely reflect differential vulnerability to persistent unfavourable habitats, as taxa with terrestrial adult stages can escape adverse aquatic conditions. Moreover, many taxa exhibiting strong negative responses (e.g., Tabanidae, *Paroxyethira* spp., *Berosus* spp., and many gastropod and worm-like organisms) are commonly associated with fine sediment and aquatic vegetation (Clapcott et al., 2017)—conditions that typically develop during prolonged low-flow periods (Rolls et al., 2012). Sediment deposition and vegetation growth—known drivers of macroinvertebrate community assembly (McKenzie et al., 2025)—can vary throughout the year, in part due to changes in river flow (Wohl et al., 2015). Future work could therefore examine how changes in sediment deposition and river-bed morphology mediate the ecological effects of flow fluctuations at different frequencies.

Fast fluctuations consistently drove unidirectional changes in community structure by reducing abundance across most taxa groups. The time window for fast fluctuations in our models captures relatively recent floods that can physically displace individuals and after which the community is unlikely to be fully recovered (Death, 2008). Expected community changes in response to fast flow fluctuations were consequently driven by abundance gradients. Yet, while most taxa experienced reduced abundance in response to fast increases in flow, the severity of impact partly depended on mobility traits. Swimmers showed strong declines, possibly because they rely on active locomotion to maintain position and must recolonise upstream habitats after being swept downstream. By contrast, burrowing or substrate-attached taxa may suffer flood mortality but still retain individuals locally (Bae & Park, 2016). Our results align with previous research showing that fast increases in flow reduce abundance across most taxa simultaneously, rather than causing shifts in taxonomic dominance (Le et al., 2021).

In addition to slow and fast flow fluctuations, seasonal fluctuations also played a notable role in shaping species composition in these dynamic ecosystems. Unlike other sources of environmental variation, the periodicity of seasonal fluctuations allows organisms to track flow variability within the course of their life cycles (Bernhardt et al., 2020; Lytle & Poff, 2004; Varpe, 2017) and can facilitate the time-sharing of a given habitat by different species (Bogan & Lytle, 2007; Tonkin et al., 2017). The community response to seasonal fluctuations could therefore be distinct from the response to other, more unpredictable fluctuations of similar frequency in many systems. Our results agree with this expectation, showing that community changes due to seasonal fluctuations were driven almost exclusively by balanced abundance changes (i.e., by the replacement of individuals of some species with individuals of others), matching previous findings (Tonkin et al., 2017). Of course, our dataset is mainly sampled at the annual frequency, with most samples taken during the austral summer, which makes the interpretation of the seasonal signal challenging. A more accurate representation of community responses to seasonal fluctuations therefore requires a higher temporal sampling resolution, including samples from seasons with different environmental conditions.

In our analyses, we have deliberately assumed linear and additive effects across different components of environmental variation, but nonlinear responses and more complex interactions are possible. Fast fluctuations superimposed on long-term directional trends could produce compounding ecological impacts. Similarly, fast fluctuations on top of seasonality could push seasonal regimes into extreme conditions. The consequences of such events are only just beginning to be understood (Bastos et al., 2023; Mahecha et al., 2022; Tonkin et al., 2026; Zscheischler et al., 2020), but high-resolution environmental datasets offer opportunities to further explore these complex dynamics. Finally, there are many possibilities with regard to choosing time windows for fast and slow fluctuations. We chose values based on prior knowledge of New Zealand rivers, but the process could be automated using model selection criteria or more complex models that optimise the time windows for each component as model parameters. The latter approach would also allow for taxa-specific time windows, which could reveal patterns in species’ responses and potential eco-evolutionary trade-offs. The potential applications of these approaches represent an interesting and challenging research agenda.

Our results strongly support the decomposition of temporal fluctuations to describe community responses to key environmental drivers, and demonstrate a methodological framework that can be readily applied across diverse ecosystems and environmental variables. Overlooking component-specific responses risks missing key mechanisms of community assembly in dynamic systems. Consequently, characterising community responses based on raw or aggregated values of environmental variables may lead to incomplete predictions of how communities respond to complex environmental change. Understanding the role of environmental fluctuations in determining species responses and community patterns is critical for predicting the consequences of human impacts, including changes in land use, water management, and climate change (Rolls et al., 2018;

Tonkin et al., 2019). However, the relative importance of different temporal components likely varies among systems. Ecosystems dominated by long-lived species may respond more strongly to slow fluctuations, while highly seasonal systems may be primarily structured by seasonal components (Hernández-Carrasco et al., 2025; Lytle & Poff, 2004; Petchey, 2000). Future research applying this framework across diverse systems will help identify the ecological, evolutionary, and environmental factors that determine which temporal frequencies are most important for community dynamics, ultimately improving our ability to predict community responses to environmental change across different contexts.

## Data and code availability

Macroinvertebrate data were derived from the National River Water Quality Network (NRWQN) monitoring programme operated by the National Institute of Water and Atmospheric Research (NIWA; now part of Earth Sciences New Zealand). River flow data were obtained from from Earth Sciences New Zealand Hydro Web Portal (hydrowebportal.niwa.co.nz). All data and code required to reproduce the results presented in this study are archived in a public GitHub repository (github.com/tonkinlab/flow freq resp) and a permanent Zenodo archive (doi.org/10.5281/zenodo.19047630).

## Author contribution

**Conceptualization:** D.H.-C., G.K., A.J.G., R.S.L., N.R.H., H.A.L.H., C.M., L.W., I.S., J.D.T.

**Methodology:** D.H.-C., G.K., A.J.G., I.S., H.R.L., J.D.T.

**Data curation:** A.J.G., R.S.L., H.A.L.H., L.W.

**Writing (original draft):** D.H.-C., G.K., A.J.G., N.R.H., H.A.L.H., J.D.T.

**Writing (review and editing):** All authors

**Visualization:** D.H.-C., A.J.G., L.W., J.D.T.

**Supervision and funding acquisition:** J.D.T.

## Acknowledgments

This work stemmed from a workshop held at the University of Canterbury’s Cass Field Station. We thank Ignacio Reyes Sainz, Anne M. McLeod, Julian Merder, and the members of the Freshwater Ecology Research Group (FERG) for discussions and feedback. This research was funded by a Rutherford Discovery Fellowship administered by the Royal Society Te Apārangi (RDF-18-UOC-007) awarded to JDT. JDT is also supported by Te Pūnaha Matatini, a Centre of Research Excellence funded by the Tertiary Education Commission, New Zealand. HRL is supported by the Marsden Fund managed by the Royal Society Te Apārangi (grant MFP-UOC2102) and the Bioprotection Aotearoa Centre of Research Excellence.

