## Supporting Information for "Population and community responses to the fast, slow, and seasonal components of environmental variation"

#### S1 Data coverage

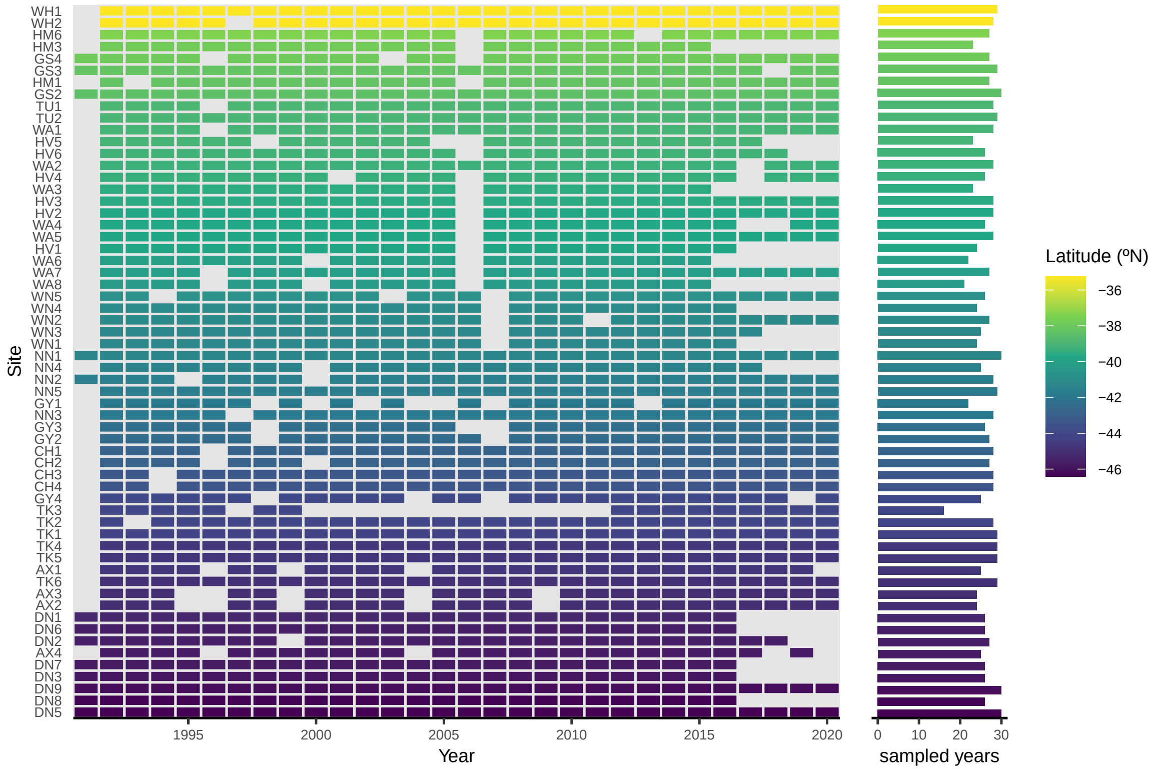


Fig. S1: **Spatial and temporal coverage of the macroinvertebrate sampling scheme.** Coloured tiles indicate years in which the river community was sampled.

#### S2 Correlation between river size and river order

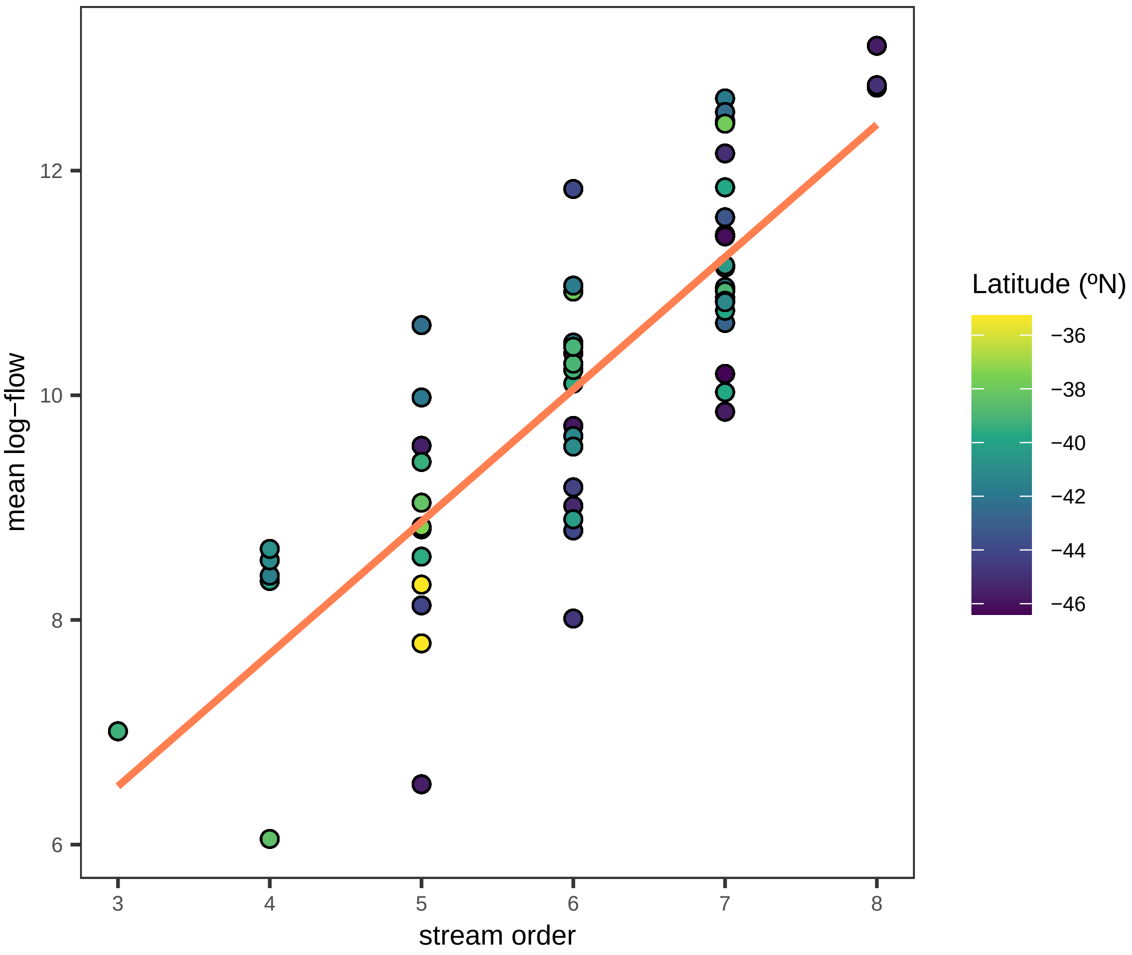


Fig. S2: **Correlation between average log-flow and stream order.**

### S3 Correlation between taxa responses to flow components

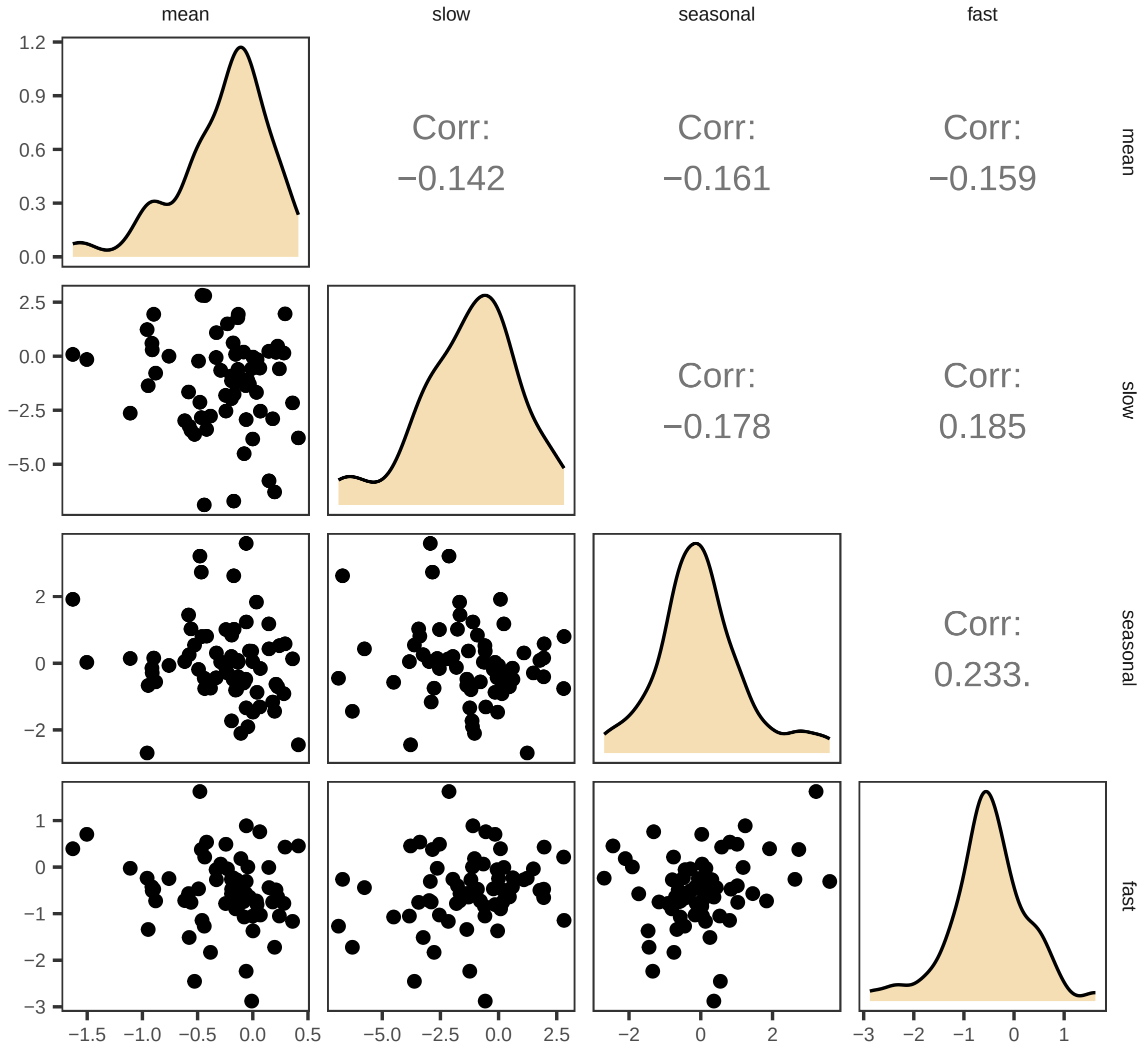


Fig. S3: **Correlation between taxon responses to flow components**. The lower triangle shows scatterplots of taxon responses, diagonal panels show the distribution of responses to each variable, and the upper triangle indicates Pearson's correlation coefficients; no significant associations were found except for a marginally significant correlation (<0.1) between the effects of fast and seasonal fluctuations on taxon abundances.

### S4 Human impact, river size, and residual variability in community composition

Our models included effects of mean flow (strongly correlated with river size; see Fig. S2) and TNER (a proxy of human impact) to account for known gradients in community composition driven by land use. TNER values were higher for sites that were classified as potentially impacted by human activity at the start of the monitoring program (Fig. S4). Taxonomic groups associated with low mean log-flow included most Ephemeroptera, Plecoptera, and Trichoptera—indicator groups of good water quality—while tolerant species such as Tanytarsini and Ceratopogonidae presented positive coefficients (Fig. S5). Ephemeroptera, Plecoptera, and Trichoptera were also generally negatively affected by TNER, although this relationship was less consistent across taxa. Taxon loadings in the two latent variables included in the model revealed a gradient from sites with taxa commonly associated with poor water quality (negative scores for LV2) to sites with more sensitive taxa (positive scores for LV2) (Fig. S6B, Fig. S7B). Site scores did not show any clear latitudinal pattern (Fig. S6A, S7A). This pattern suggests that the two latent variables mainly accounted for additional variability in river water quality.

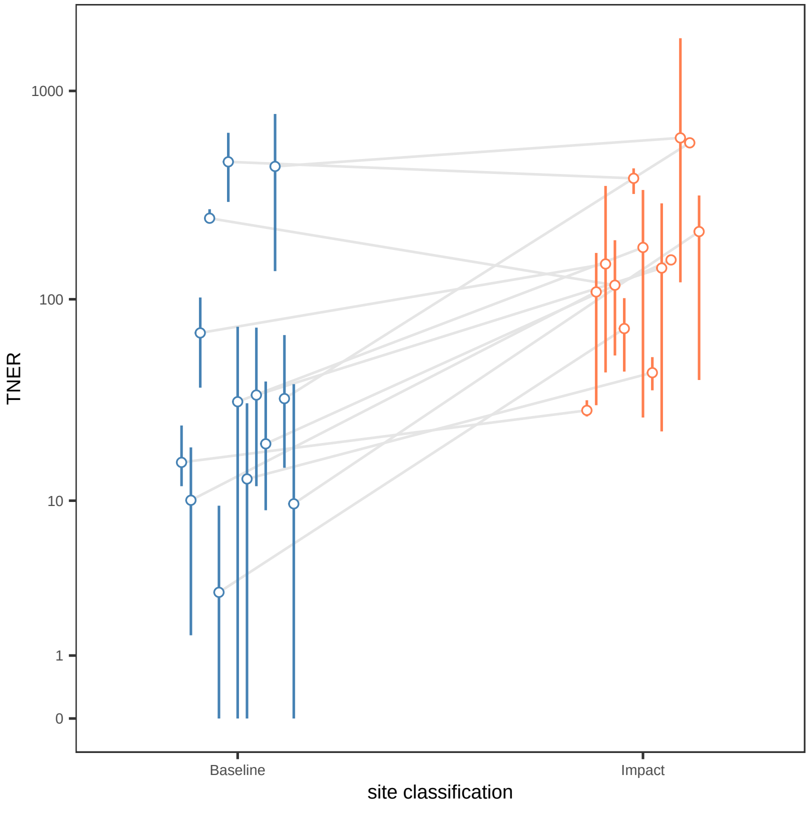


Fig. S4: **Comparison of Total Nitrogen Excretion Rates (TNER) between streams considered *impacted* and *baseline* at the start of the monitoring program.** Line ranges indicate the temporal variability in each river.

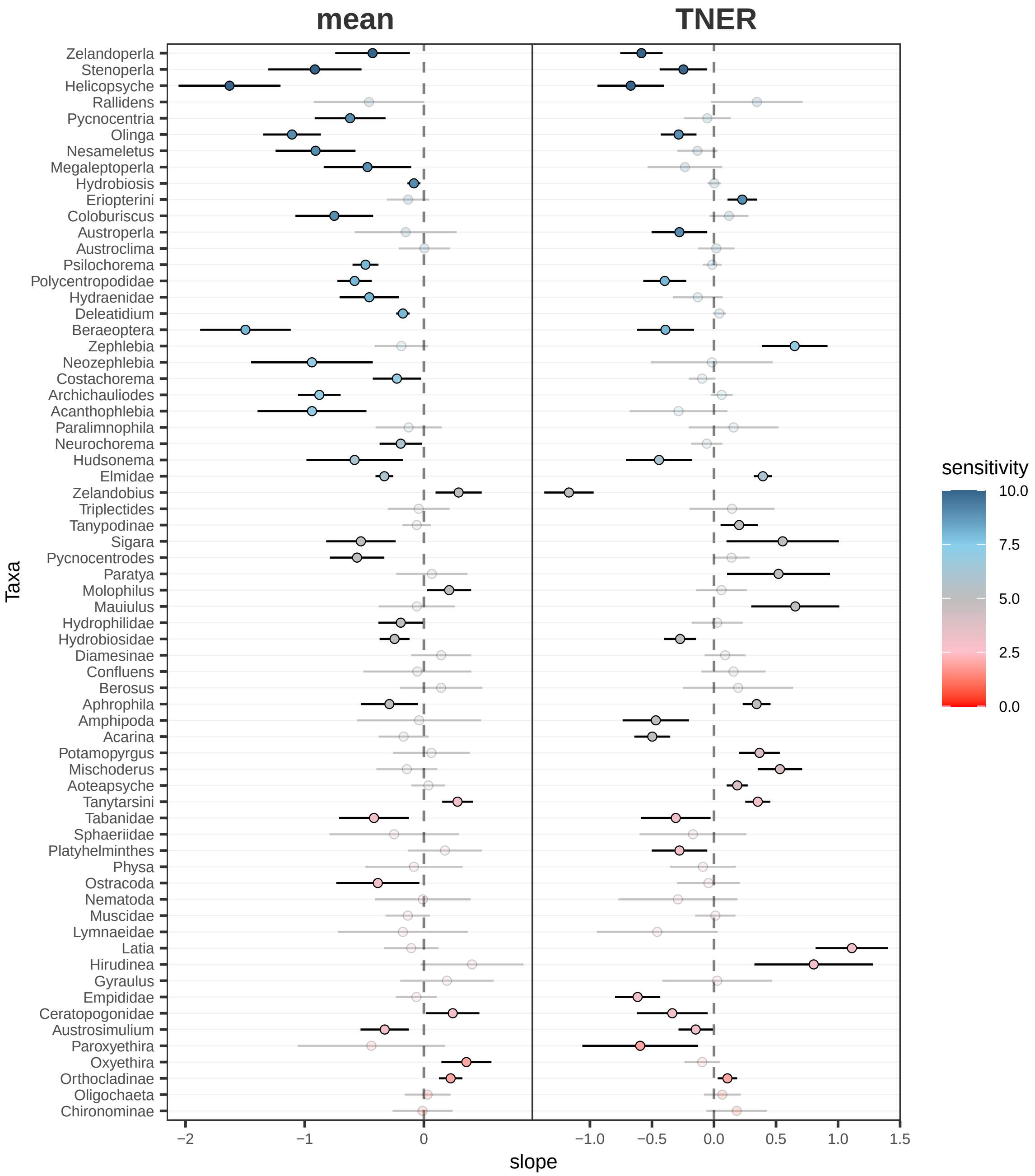


Fig. S5: **Macroinvertebrate responses to mean flow and TNER.** Colour gradient indicates the sensitivity score of each taxon group.

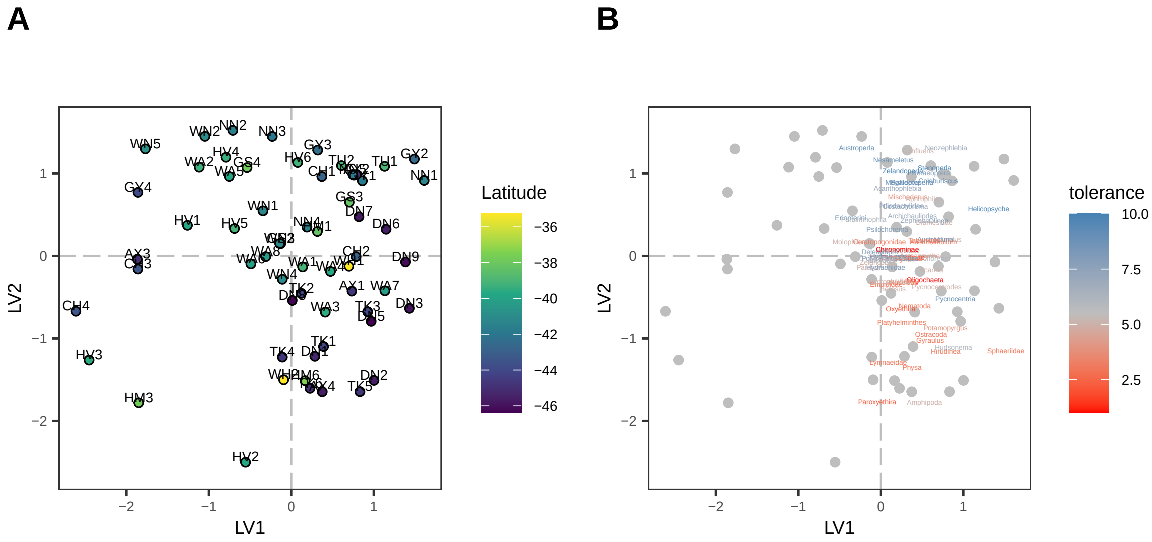


Fig. S6: **Model-based ordination of site loadings without traits.** Species loadings have been re-scaled by a factor of 0.6 for representation purposes.

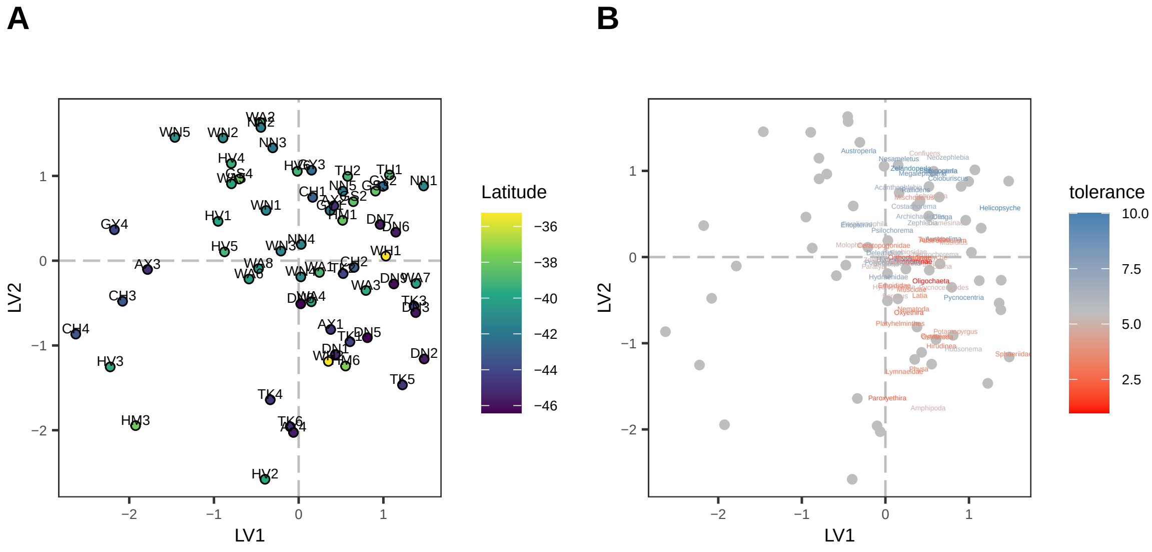


Fig. S7: **Model-based ordination of site loadings with traits.** Species loadings have been re-scaled by a factor of 0.6 for representation purposes.
